# Modulation of SARS-CoV-2 Spike-induced Unfolded Protein Response (UPR) in HEK293T cells by selected small chemical molecules

**DOI:** 10.1101/2021.02.04.429769

**Authors:** B Balakrishnan, K Lai

## Abstract

Coronaviruses (CoV) exploits the endoplasmic reticulum (ER) of the host cells for replication and in doing so, increases ER stress. evokes Unfolded Protein Response (UPR) and possibly autophagy, which could all attribute to the pathophysiology of the viral infections. To date, little is known about the roles of ER stress, UPR, and autophagy in SARS-CoV-2 infection. Here we over-expressed the viral Spike (S) protein in cultured HEK293T cells, as it has been shown that such protein is largely responsible for UPR activation in other CoV-infected cells. We noticed, in the transfected cells, heightened ER stress, activation of the PERK-eIF2α arm of the UPR, induction of autophagy and cell death. When we treated the transfected cells with Tauroursodeoxycholic acid (TUDCA), 4-phenyl butyric acid (PBA), Salubrinal, Trazadone hydrochloride, and Dibenzoylmethane (DBM), we saw reduced the BiP/GRP78 levels, but only PBA and TUDCA could significantly diminish the levels of peIF2α and autophagy expression.

## 1. Introduction

Coronavirus disease 2019 (COVID-19) is a potentially lethal, debilitating infectious disease caused by the newly discovered coronavirus called SARS-CoV-2 [1]. Like other coronaviruses, SARS-CoV-2 is an enveloped and positive-sense single-stranded RNA virus with a large genome of ~30 kb. Morphologically, coronaviruses are spherical or pleomorphic in shape with a mean diameter of 80–120 nm [2]. They are characterized by the large (20 nm) “club-like” projections on the surface, which are the heavily glycosylated trimeric Spike (S) proteins [2, 3]. Coronavirus infection is initiated by the binding of the S protein to the angiotensin-converting enzyme 2 (ACE2) cell surface receptors followed by fusion of the virus and cell membrane to release the viral genome into the cell [4, 5]. The S protein comprises two functional subunits, S1 (bulb) for receptor binding and S2 (stalk) for membrane fusion. The interaction between the host ACE2 cell surface receptor and the S1 subunit is the central element of coronaviruses’ tropism [6]. It replicates in the cytoplasm, and its life cycle is closely associated with the endoplasmic reticulum (ER). The viral activities thus have a profound impact on ER functions. Particularly, SARS-CoV-2 hijacks the ER to process its structural and nonstructural proteins.

The accumulation of nascent and unfolded viral secretory and structural proteins in the ER lumen can lead to ER stress and the downstream activation of multiple signaling pathways [7]. To accommodate the biosynthetic encumbrance and capacity of the ER for maintaining cellular homeostasis, the host cell activates Unfolded Protein Response (UPR) to mitigate the ER stress. The UPR comprises three critical signaling pathways initiated by localized stress sensors in ER, such as inositol-requiring enzyme 1 (IRE-1), activating transcription factor 6 (ATF6), and PKR-like ER kinase (PERK) [8]. These proteins send robust signals to the cytosol and nucleus to alleviate the burden of misfolded protein and ensure successful ER protein homeostasis [9]. However, undue or prolonged activation of UPR can also trigger autophagy and apoptosis [10], which could play a significant role in the pathophysiology of viral infections. For SARS-CoV (the coronavirus that caused SARS), it was reported that the Spike (S) protein, but not the envelope (E), membrane (M), or the nucleocapsid (N) protein evokes UPR by the transcriptional activation of GRP78 (BiP)/GRP94 and upregulation of the PERK pathway with no effects on the other two arms of UPR [11–13]. Studies also confirmed UPR-triggered autophagy of the host cells in SARS-COV and MERS-COV infections [14, 15]. Based on the previous studies on other coronaviruses, it is reasonable to assume that SARS-CoV-2 may induce autophagy *via* UPR activation in the infected cells. However, emerging findings suggest that SARS-CoV and COVID-19 are also unique in many ways. For instance, short-term loss of smell and taste during the infection cycle [16–19], abnormal coagulation [20–22], silent hypoxia [23], and rash [24–26] in some patients with the new COVID-19 are just some of the *unique* phenotypes that were rarely reported for other coronaviral illnesses. Such unique phenotypes are telltale signs that it is premature to assume everything we learned from other coronaviruses, including SARS-CoV, can be applied in wholesale to SARS-CoV-2 without further investigations.

In this short communication, we aim to study the impact of the over-expression of the Spike (S) protein of SARS-CoV-2 in HEK293T cells on ER stress manifestation and its downstream effects, including autophagy. Puelles and coworkers recently reported the SARS-CoV-2 tropism in human kidney by detecting the presence of ACE2 mRNA in the kidney cells, as well as the enhanced expression of transmembrane serine protease 2 (TMPRSS2) and cathepsin L (CTSL), which are also considered to facilitate SARS-CoV-2 infection in multiple kidney-cell types from fetal development through adulthood [27]. Indeed, acute kidney injury and dysfunctions are observed in a large proportion of COVID-19 patients [28–31]. Hence the use of HEK293T cells in this study is of significant clinical relevance. Our results showed that over-expression of the viral Spike protein in the HEK293T cells activated UPR, which led to autophagy. Interestingly, the activated UPR and the downstream autophagy induced could be substantially diminished by selected UPR modulators, some approved by the U.S. Food and Drug Administration (FDA) for treating other diseases [32–35].

## 2. Materials and Methods

### 2.1. Cells, Antibodies, and Reagents

HEK293T cells obtained from ATCC (Manassas, VA) were cultured in a 37°C, 5% CO_2_ incubator in Dulbecco’s Modified Eagle Medium (DMEM) supplemented with 10% (vol/vol) FBS, 10 mM HEPES, 100 IU/mL penicillin, and 100 μg/mL streptomycin.

Anti-GRP78 (BiP), anti-peIF2α, anti-LC3B, and anti-GAPDH were purchased from Cell Signaling Technology Inc. (Danvers, MA). Goat anti-rabbit IRDYE 800CW was purchased from Li-COR Biosciences. (Lincoln, NE). anti-SARS-CoV-2 Spike Protein was purchased from Origene Technologies Inc. (Rockville, MD).

Transfection reagent Lipofectamine^®^ 3000 transfection reagent was purchased from Thermo Fisher Scientific Inc. (Waltham, MA).

Cell Proliferation Kit I (MTT) was obtained from Sigma-Aldrich Inc. (St. Louis, MO).

Tauroursodeoxycholic acid (TUDCA), 4-phenyl butyric acid (PBA), Salubrinal, Trazadone hydrochloride, and Dibenzoylmethane (DBM) were purchased from Sigma-Aldrich Inc. (St. Louis, MO).

### 2.2. Plasmid

pcDNA3.1-SARS2-Spike, which was constructed by Fang Li (Addgene plasmid # 145032; http://n2t.net/addgene:145032; RRID: Addgene_145032), was purchased from Addgene Inc. (Watertown, MA).

### 2.3. Transfection Studies

HEK293T cells were plated onto six-well plates at a density of 4×10^5^ cells per well and cultured at 37°C with 5% CO_2_ overnight for transfection. A total amount of 5 μg DNA per well was used for transient transfection with Lipofectamine^®^ 3000 according to the manufacturer’s protocol. Twenty-four hours post-transfection, culture media was changed, and the cells were extracted and lysed using MPER lysis buffer (Thermo Fisher Scientific Inc.). For the drug treatment, the UPR modulators at the indicated concentrations were administered 4 hours post-transfection, and proteins from the cells were extracted after 24 hours of treatment.

### 2.4. Western Blot Analysis

Cells overexpressing C9-tagged Spike protein and the drug treatment group were harvested 24- and 28-hours post-transfection respectively. The cells were lysed with 0.1 ml of MPER buffer containing 1 mM NaVO3, 2.5 mM sodium pyrophosphate, 1 mM β-glycerol phosphate, 1 mM phenylmethylsulfonyl fluoride, 5 μg/ml aprotinin, and 5 μg/ml leupeptin, (pH 7.5). 30 μg of total protein from cell lysate was separated on 12% sodium dodecyl sulfate-polyacrylamide gel electrophoresis, electro-transferred to nitrocellulose membrane (LI-COR). Antibodies described above were used to probe the membranes according to the protocols the manufacturer provided. The signals were detected by Infrared detection using an Odyssey scanner and analyzed by Image Studio software (LI-COR).

### 2.5. Detection of Autophagy

For the microtubule-associated protein 1A/1B-light chain 3 (LC3) mobility shift assay, SARS-CoV-2 Spike protein-expressing cells were washed with cold PBS, lysed with MPER buffer, and subjected to Western blot analysis with antibodies against LC3B. LC3-I is about 16 kD, and lipidated LC3 (LC3-II) is about 14 kD. The level of LC3 II formation represents the autophagic activity [36].

### 2.6. Cell viability determination by MTT assay

Cell viability was assessed by an MTT assay. Briefly, HEK293T cells were seeded in 96-well plates at a density of 5×10^4^/well. The cells were transfected with the plasmid, pcDNA3.1-SARS2-Spike as described above. 24 hours post-transfection, MTT solution (10 μl, 5 mg/ml in PBS) was added to each well and incubated at 37°C for 4 hours. Then, the medium was replaced by 100 μl DMSO per well. The plate was gently shaken for 5 minutes to completely dissolve the precipitate and incubated for 30 minutes. The absorbance was measured at 560 nm using a microplate reader. Cell viability was expressed as a percentage of the control.

### 2.7. Quantification and Statistical Analysis

All data were expressed as Mean± SEM as indicated. Statistical significance between the two groups was tested by the Student’s t-test. P values of less than 0.05 were considered significant.

## 3. Results

### 3.1. UPR is activated upon SARS-CoV-2 Spike protein overexpression in cultured cells

To study the molecular events subsequent to SARS-CoV-2 Spike overexpression in details, the plasmid expressing SARS-CoV-2 Spike protein was transiently transfected into cultured HEK293T cells at different DNA concentrations. Manifestation of ER stress was first demonstrated by evaluating the intracellular GRP78 levels in the transfected cells. In cells over-expressing the SARS-CoV-2 Spike protein, the level of BiP/GRP78 was significantly higher compared to the untransfected control (Fig. 1a). We next investigated which specific UPR pathways are activated upon overexpression of the Spike protein, with particular emphasis on the PERK arm. We found a significant increase in the levels of phosphorylated eIF2α in the transfected cells 24 hours post-transfection (Fig. 1b). This further substantiated the hypothesis that SARS-CoV-2-Spike protein overexpression induces UPR in cultured cells.

**Figure 1.**
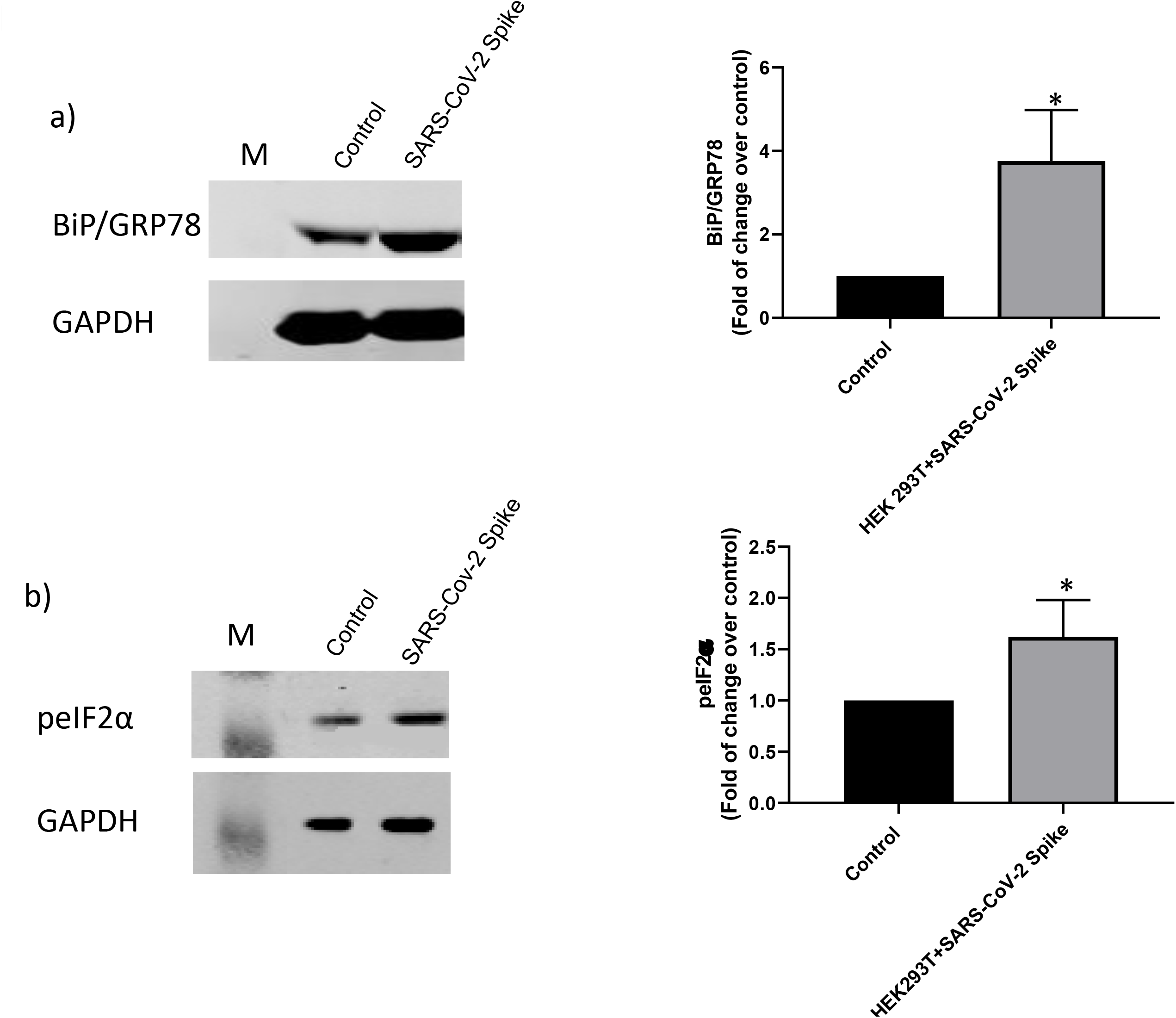
Upregulation of BiP/GRP78 in HEK293T cells over-expressing the SARS-CoV-2 Spike protein. HEK293T cells were transfected with pcDNA3.1-SARS2-Spike and the protein levels of (a) BiP/GRP78 and (b) phosphorylated eIF2α (peIF2α) were determined by Western blotting using specific antibodies 24 hours after transfection. Quantified values of the specific protein abundance, which were normalized to the abundance of GAPDH, were included at the right panel.

### 3.2. SARS-CoV-2 Spike protein expression triggers autophagy response in cultured cells

There is growing evidence to show that various host cellular responses, including autophagy, innate immunity, and apoptosis, are affected or activated by coronaviruses infection *in vivo* [37–39]. Among the many signaling pathways, the UPR and autophagy are tightly interconnected and were shown to be essential for viral infection by multiple previous studies. The implication of autophagy in coronavirus infection has attracted substantial attention, due to the SARS-CoV outbreak in 2002-2003 [40–43]. To investigate the involvement of autophagy in cells over-expressing the SARS-CoV-2 Spike protein, we evaluated the molecular markers of autophagy by monitoring levels of LC3-I and LC3-II. We found that levels of LC3-II were increased in Spike protein-overexpressed cells from 24 hours post-transfection (Fig. 2a), which is indicative for an activated autophagy response that culminates in degradation cytoplasmic components in the lysosomes. Collectively, these data illustrated that the PERK/eIF2α pathway of UPR is activated in SARS-CoV-2 infection, along with the induction of autophagy. To assess the cell viability by Spike protein overexpression, we conducted MTT assay 24 hours post transfection in HEK293T cells. The results revealed that SARS-CoV-2 Spike protein expression in HEK293T cells leads to cell death (Fig. 2b), which could be the consequence of UPR-induced autophagy shown above.

**Figure 2.**
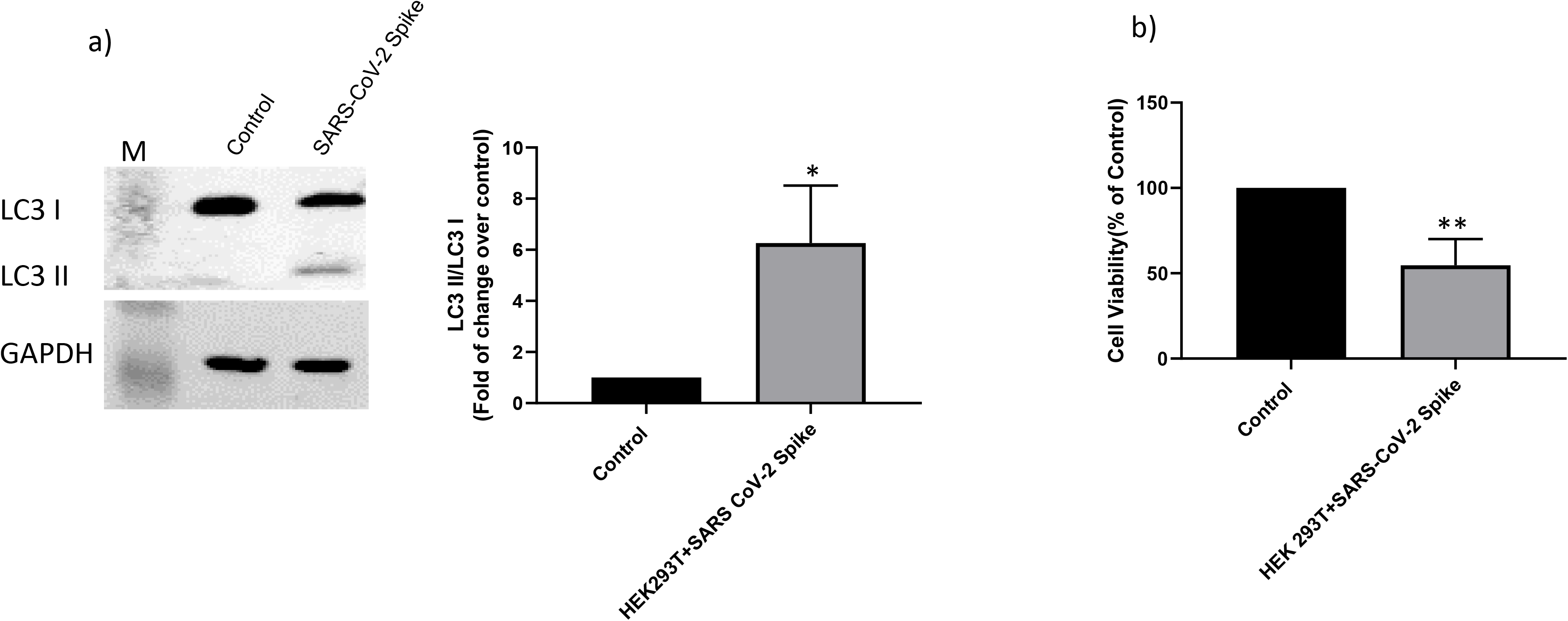
SARS-CoV-2 Spike protein over-expression triggers autophagy through UPR. (a) HEK293T cells were transfected with pcDNA3.1-SARS2-Spike and the protein levels LC3I and LC3-II were determined by Western blotting using anti-LC3B antibody after 24 hours. Values were normalized to GAPDH. The ratio of LC3-II over GAPDH is graphically represented on right. (b) Cell viability of the transfected cells was evaluated by MTT assay and the quantified values were normalized to non-transfected controls.

### 3.3. UPR modulators reduced the UPR and autophagy in cultured cells

We next selected a few UPR modulators, some of them were approved by the FDA to treat other diseases [32–35], in an attempt to modulate the level or ER stress and/or the UPR in HEK293T cells over-expressing the viral Spike protein. HEK293T cells were treated with 4-PBA, TUDCA, DBM, Trazadone, and Salubrinal at indicated concentrations 4 hours after plasmid encoding Spike protein transfection. The expressions of GRP78, peIF2α, and LC3-II were examined at 24 hours post-transfection. Salubrinal, TUDCA, PBA, and Trazadone significantly reduced the BIP/GRP78 levels in cells overexpressed with viral Spike protein (Fig. 3a). In addition, we saw a significant reduction in peIF2α in cells treated with the PBA and TUDCA (Fig. 3b). Interestingly, treatment with PBA and TUDCA also resulted in a diminished level of the autophagy marker, LC3-II (14 kD) in the cells over-expressing Spike at 24 hours (Fig. 3c). Altogether, we demonstrated the cause-and-effect relationship between Spike protein-induced UPR and the subsequent autophagy. We further analyzed the cell survival with UPR modulators and the data confirmed the protective effects of TUDCA and PBA on cell survival (data not shown).

**Figure 3.**
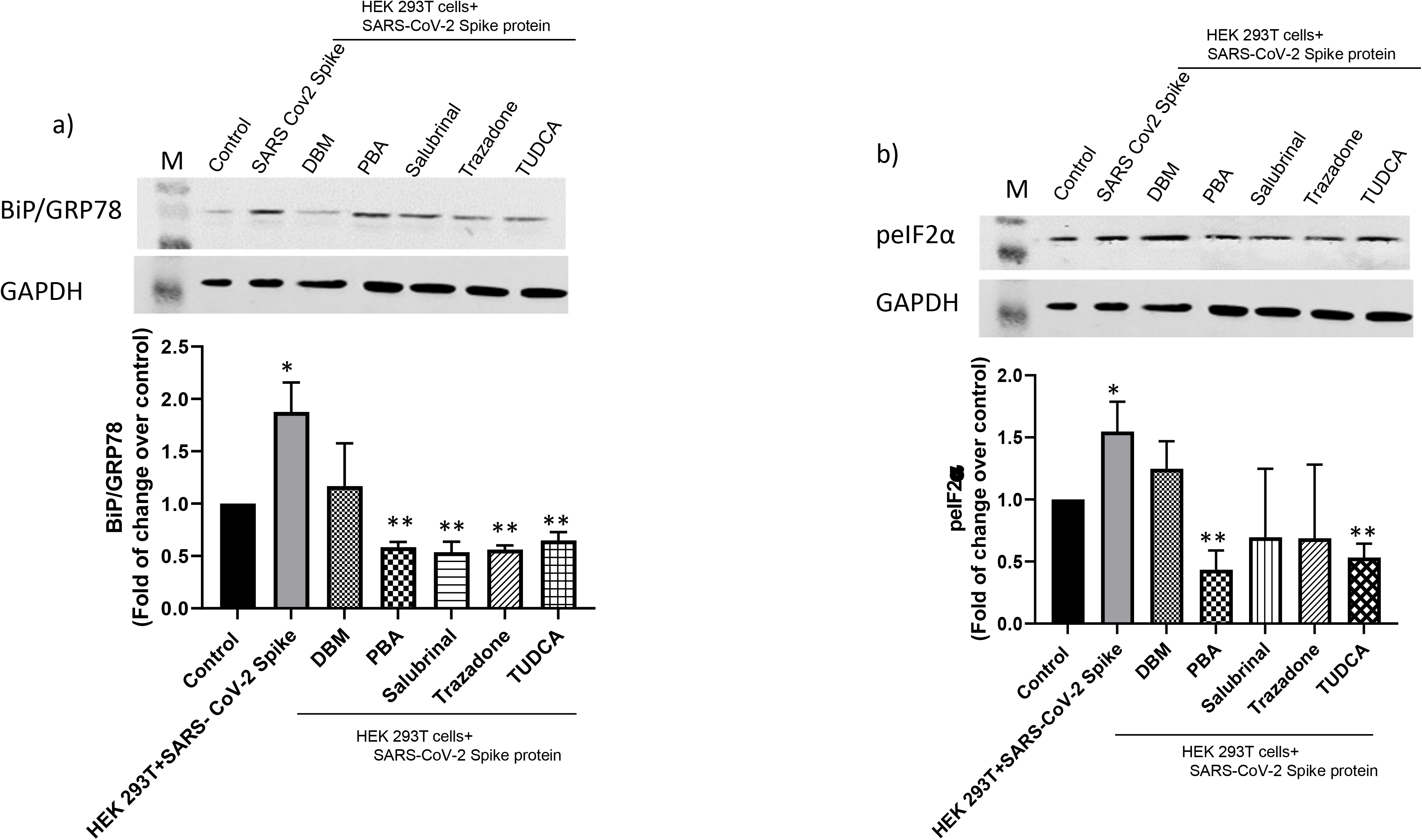
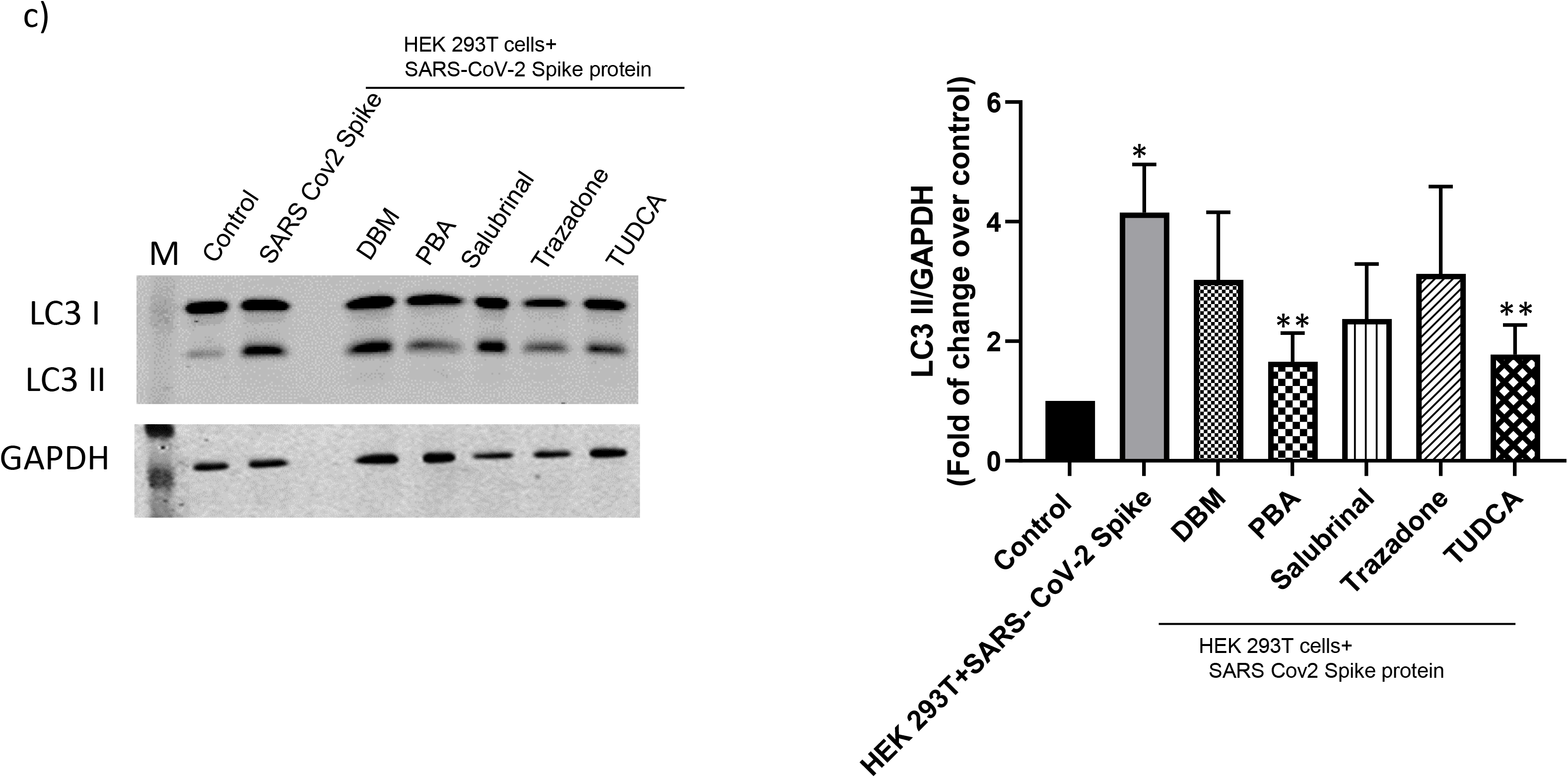
UPR modulator treatment reduced levels of ER stress, UPR and autophagy. HEK293T cells were transfected with pcDNA3.1-SARS2-Spike and 4 hours after transfection, the cells were treated with the selected UPR modulators, Salubrinal (50μM), 4-PBA (2mM), TUDCA (100μM), DBM (50μM) and Trazadone (50μM) for 24 hours. The protein levels of (a) BiP/GRP78, (b) phosphorylated eIF2α were determined by Western blotting using specific antibodies. Values were normalized to that abundance of GAPDH and were presented in the graphs on the right. (c) The protein levels LC3-II with UPR modulator treatment were determined by Western blotting and normalized to the abundance of GAPDH. The ratio of LC3 II over GAPDH is graphically represented on right. * represents significant values compared to control. ** represents significant values compared to HEK293T + SARS-CoV-2 Spike.

## 4. Discussion

Since February 2020, our world has been embroiled in a pandemic that is unprecedent in scale. As of Dec 27^th^, 2020, there are more than 81 million people worldwide infected with SARS-CoV-2_and over 1.77 million deaths. This horrific loss of human lives and the emotional toll inflicted on the loved ones of the deceased will continue to rise until the spread of the disease is halted. At the same time, many survivors who were fortunate to escape death suffer from a host of complications such lung fibrosis or systemic organ damages that will take years to fully heal. In addition to the significant mortality and morbidity caused by COVID-19, millions of jobs have been lost in the U.S. alone since March. Therefore, there is an urgent need for the development of life-saving therapies and preventive measures for COVID-19. But to accomplish these goals, we must improve our understanding of the pathophysiological mechanisms of the disease.

Being a coronavirus, SARS-CoV-2 shares some similarities with other Coronaviruses, including SARS-CoV. While our current knowledge on existing human coronaviral diseases could offer important insights into SARS-CoV, emerging findings suggest that SARS-CoV-2 is also unique in many ways. Some of the phenotypes such as short-term loss of smell and taste [16–19], abnormal coagulation [20–22] seen in many patients with the new COVID-19 were rarely reported for other coronaviral illnesses. Therefore, it is premature to assume the knowledge we gained from the studies of other coronaviruses, including SARS-CoV, can be applied in entirety to SARS-CoV-2 without additional research.

In this study we focused on the potential roles of ER stress, UPR and autophagy in SARS-CoV-2 infection because such fundamental cellular processes have been shown to contribute to the pathophysiology of other diseases [44–47]. Also, in case of viral infections, the role of ER during viral replication is well-documented, and the activation of UPR has been reported in other coronaviruses-infected cells [48–51]. Therefore, the induction of ER stress and UPR activation is a crucial factor in virus-host interaction and can significantly influence the patient’s antiviral response and [52–55].

To study the potential roles of UPR and autophagy in SARS-CoV-2 infections, we chose to overexpress the SARS-CoV-2 Spike (S) protein in cultured HEK293T cells because it has been shown in SAR-CoV that the S protein is the viral protein largely responsible for activation of UPR [11–13]. Moreover, our focus on the HEK293T cells is clinically relevant due to the high evidence of acute kidney injury in patients with COVID-19. The mechanism(s) of renal injury caused by SARS-CoV-2 has not yet been fully elucidated, although ER stress/UPR has been implicated in renal injury induced by ischemia-reperfusion and nephrotoxicity [56].

Here we showed that the SARS-CoV-2 Spike protein expression in the HEK293T cells led to up-regulation of BiP/GRP78 (Fig. 1) and the activation of the PERK branch of the UPR pathway. These findings corroborated with the positive detection of ER stress markers in the lung samples infected from COVID-19 patients with severe pneumonia [57]. Our data also agreed with the recent study that shows the SARS-CoV and MHV-CoV infections activated PERK arm of UPR with subsequently increased phosphorylation of eIF2α in the host cells [58–60]. It should be noted that the PERK pathway regulates innate immunity by suppressing type 1 interferon (IFN) signaling [60]. Hence, it suggests that UPR does play a role in attenuating IFN responses and innate immunity in coronavirus-infected cells.

Several lines of evidence demonstrated that ER stress and the associated UPR contributes to autophagy [61, 62]. However, the specific UPR pathway activated to mediate autophagic activities, and whether cytoprotective autophagy or autophagic cell death is induced upon ER stress, is not entirely clear and probably varies with the different triggers of the UPR [63, 64]. Nonetheless, an essential step in UPR-induced autophagosome formation is the phosphorylation of PERK/eIF2α [65]. Once phosphorylated, eIF2α can induce the production of LC3-II from LC3-I to induce autophagy [66]. To further investigate the potential roles of UPR and autophagy in SARS-CoV-2, we examined the HEK293T cells over-expressing the Spike protein and found that the LC3-II protein level was significantly upregulated with Spike protein expression, leading to cell death (Fig. 2). Furthermore, the changes in LC3-II observed were significantly reversed by pretreatment with PBA and TUDCA (Fig. 3), indicating these compounds could block the autophagy activation induced by over-expression of the Spike protein. It is unclear why PBA and TUDCA are more effective than the other UPR modulators we selected in this study. However, PBA and TUDCA are chemical chaperones [32, 67–71] and they could act to minimize UPR induction earlier on rather than mitigating the downstream signaling pathway like Salubrinal and the others.

At present, clinical management of COVID-19 includes infection prevention (mask-wearing, social distancing, vaccination)supportive care (supplemental oxygen and mechanical ventilatory support) and limited and selective use of new anti-viral therapeutics [Remdesivir (Veklury) [72], convalescent plasma therapy with mixed results [73]. Dexamethasone, a glucocorticoid used to reduce inflammations in patient cells, has shown some promises in reducing the risk of death for patients with more severe SARS-CoV-2 infections, although the molecular mechanisms involved remain unclear. However, it has been shown that such glucocorticoid ameliorates ER stress in intestinal secretory cells with protein misfolding *in vitro* and *in vivo* [74]. Therefore, our current studies support the hypothesis that Dexamethasone might act through the modulation of UPR in the infected cells of the COVID-19 patients. Given the potential side-effects of prolonged use of glucocorticoids [75], one might want to consider the alternate use of other UPR modulators such as PBA and TUDCA, in combination of other approved antivirals.

## 5. Conclusions

We show for the first time that over-expression of the Spike (S) protein of SARS-CoV-2 heightened ER Stress, activated UPR and caused autophagy in human HEK293T cells in culture. Such cellular responses were significantly diminished by specific chemical chaperones PBA and TUDCA, but not other UPR modulators. These findings render significant implications in the pathophysiology and the treatment of the viral infection.

## Notes

### Competing Interest Statement

The authors have declared no competing interest.

